# Human defensive freezing is associated with acute threat coping, long term hair cortisol levels and trait anxiety

**DOI:** 10.1101/554840

**Authors:** Mahur M. Hashemi, Wei Zhang, Reinoud Kaldewaij, Saskia B.J. Koch, Rosa Jonker, Bernd Figner, Floris Klumpers, Karin Roelofs

## Abstract

The detection and anticipation of threat facilitates innate defensive behaviours including freezing reactions. Freezing in humans is characterized by reductions in body sway and heart rate and limited evidence suggests that individual differences in freezing reactions are associated with hypothalamic-pituitary-adrenal (HPA) axis activity and anxiety. However, previous measurements of human freezing reactions were largely based on passive threat contexts where natural variations in adaptive threat coping could not be assessed. In a well powered sample (N=419), we studied individual differences in anticipatory freezing reactions, by measuring body sway and heart rate, during an active shooting task where shooting decisions had to be taken under threat of shock. We linked freezing measures to subsequent actions and predictors of anxiety-related psychopathology, including accumulated long-term (3 months) hair cortisol concentrations (HCC) and trait anxiety. The anticipation of threat of shock elicited significant body sway- and heart rate reductions consistent with freezing. Whereas both freezing-related reductions in body sway and heart rate were associated with faster correct shooting decisions, body sway reductions were additionally related to more impulsive shooting (false alarms). Individual differences in threat-related reductions in body sway but not heart rate were further associated to lower HCC and higher trait anxiety. The observed links between freezing and subsequent defensive actions as well as predictors of stress-related psychopathology suggest the potential value of defensive freezing reactions as somatic marker for stress-vulnerability and resilience.

## Introduction

The detection of threat elicits a range of defensive behaviours varying from hard-wired, automatic, initial reactions to subsequent learned instrumental actions (Ledoux, Daw, & Emerson, 2018). In this cascade of defensive behaviours lies a vast potential for individual variation (McNaughton & Corr, 2004; Niermann, Figner, & Roelofs, 2017). It is important to understand this variability for more than theoretical purposes, as maladaptive threat processing is key to anxiety disorders which are a burden for many individuals in our society (Lewis-Fernandez et al., 2010; Robinson, Charney, Overstreet, & Grillon, 2012).

Freezing is an innate defensive reaction -across species-that is characterized by movement cessation. Upon distal threat detection and in anticipation of potential threat, human freezing is typically accompanied by heart rate deceleration (or: bradycardia) (Azevedo et al., 2005; Facchinetti, Imbiriba, Azevedo, Vargas, & Volchan, 2006; Hagenaars, Oitzl, & Roelofs, 2014). Prolonged bradycardia responses have been suggested to enhance sensory processing to optimize the detection of threatening information in the environment (Campbell, Wood, & McBride, 1997; Graham & Clifton, 1966; Lojowska, Gladwin, Hermans, & Roelofs, 2015; Vila et al., 2007). At this stage, freezing reactions are thought to facilitate risk assessment and prepare subsequent defensive actions while minimizing the chance of being detected by predators or conspecifics (Eilam, 2005; Fanselow, 1994; Roelofs, 2017). Recent studies in humans support the view that freezing can also facilitate subsequent defensive actions as they show stronger anticipatory freezing-related reactions when subjects can perform an action to avoid threat (Gladwin, Hashemi, van Ast, & Roelofs, 2016; Löw, Weymar, & Hamm, 2015). In addition, anticipatory midbrain/periaqueductal gray activity, known to mediate freezing responses in rodents (LeDoux, Iwata, Cicchetti, & Reis, 1988), predicted faster subsequent defensive actions in humans (Hashemi et al. in press). This suggests that while freezing is often considered a purely passive reaction, it may involve processes that facilitate subsequent adaptive coping actions.

Regardless of the potential role of freezing in subsequent defensive action, across species individual differences in freezing reactions have been reported as a function of biological anxiety-markers (Fox, Shelton, Oakes, Davidson, & Kalin, 2008; Frank et al., 2006; Kalin, Shelton, Rickman, & Davidson, 1998). For example, trait and state anxiety have been associated with hypervigilance and enhanced responding to threat including defensive reactions such as freezing (Frank et al., 2006; Niermann et al., 2017; Roelofs, Hagenaars, & Stins, 2010). One pathway that regulates immediate reactions to threat and may explain individual variability in freezing is the hypothalamic-pituitary-adrenal (HPA)-axis. Threat exposure rapidly activates the HPA-regulated endocrine cascade, starting with the release of corticotropin-releasing factor (CRF), which leads to the production of glucocorticoids (GCs). GCs in turn facilitate energy mobilization, recovery and normalization to homeostasis, as well as adaptation of physiological responses to future threat (De Kloet, Joëls, & Holsboer, 2005; Joels, 2017; Oitzl, Champagne, van der Veen, & de Kloet, 2010). Studies in rats and primates demonstrated the involvement of the HPA-axis in freezing reactions by showing a positive association between threat-induced freezing and increased basal cortisol levels in anxious animals (De Boer, Slangen, & Van der Gugten, 1990; Dettmer, Novak, Suomi, & Meyer, 2012; Kalin et al., 1998; Núñez, Ferré, Escorihuela, Tobeña, & Fernández-Teruel, 1996).

Limited evidence suggests that also in humans, individual differences in freezing reactions are associated with HPA-axis activity and anxiety (Niermann et al., 2017; Roelofs et al., 2010). Niermann et al. (2017), for instance, showed that prolonged freezing in adolescents with lower levels of basal salivary cortisol was linked to increased levels of internalizing symptoms (anxiety and depression). However, in the historical majority of studies of threat processing (Ledoux, Moscarello, Sears, & Campese, 2016), freezing was measured in a purely passive context, leaving open the question of whether similar associations exist under conditions of active coping. Additionally, no studies have investigated human freezing in relation to long-term accumulated cortisol concentrations. This can be reliably assessed in hair and is potentially more robustly related to stress-coping and stress-related vulnerability (for a review, see Stalder & Kirschbaum, 2012). Indeed recent work in non-human primates showed that low hair cortisol was linked to increased freezing reactions to intruder threat (Hamel et al., 2017). In humans, blunted hair cortisol concentrations (HCC) have been repeatedly linked to anxiety and stress psychopathology (Steudte et al., 2011, 2013) but relations to freezing have not yet been explored.

The aim of the current study is to assess individual differences in anticipatory freezing reactions within an active threat context and explore associations with 1) subsequent defensive action, 2) HCC and 3) trait anxiety. As research on individual differences typically requires large samples to detect reliable relationships (Hedge, Powell, & Sumner, 2017b), we tested our research question in an adequately powered sample of participants at the baseline measurement of an ongoing longitudinal study (total N=419). During an active shooting task (Gladwin et al., 2016), participants had to make timely shooting decisions under threat of shock while body sway and bradycardia were recorded as indices of anticipatory freezing reactions. We hypothesized that enhanced freezing reactions link to subsequent threat coping actions, operationalized as speed and accuracy of shooting actions under threat (Hashemi et al., in press), and are associated with long-term low HCC and high trait anxiety as indicators of anxiety-related vulnerability.

## Methods

### Participants

This cohort is part of a prospective study on the role of defensive reactions on trauma resilience in police recruits. Details on the design and methods of the full study can be found in Netherlands Trial Registry (NTR6355) and in our protocol article (Koch et al., 2017). The project was approved by the Independent Review Board Nijmegen (IRBN registration number NL48861.072.14) and was conducted in accordance with these guidelines. All participants gave written informed consent before the start of the experiments. Participants were 427 individuals (313 men and 106 women, mean age 24, with a range between 18 and 45) including 337 students from the Dutch Police Academy and 82 age-, gender-, and education-matched healthy participants from other academies, including sports academy. Eight participants were excluded from the analysis because of discontinuation of the task (n=4), hardware problems (n=1), non-compliance to task instructions (n=2), or data loss (n=1) leaving a total sample of 419 participants. At the time of the baseline measurement, police recruits were within the first year of their education, had not yet received specific shooting-related training, and performed very little active police duty. Police recruits and matched civilians were therefore not treated as different groups within this study because our hypotheses were similar for both groups at the baseline measurement. Exclusion criteria were current psychiatric and neurological disorders, a history of - or current - endocrine or neurological treatment, current drug or alcohol abuse, and current use of psychotropic medication.

### Task

The shooting task (see Figure 1) is a speeded decision-making task under threat of shock (Gladwin et al., 2016) designed to elicit anticipatory freezing reactions prior to action.

**Figure 1.**
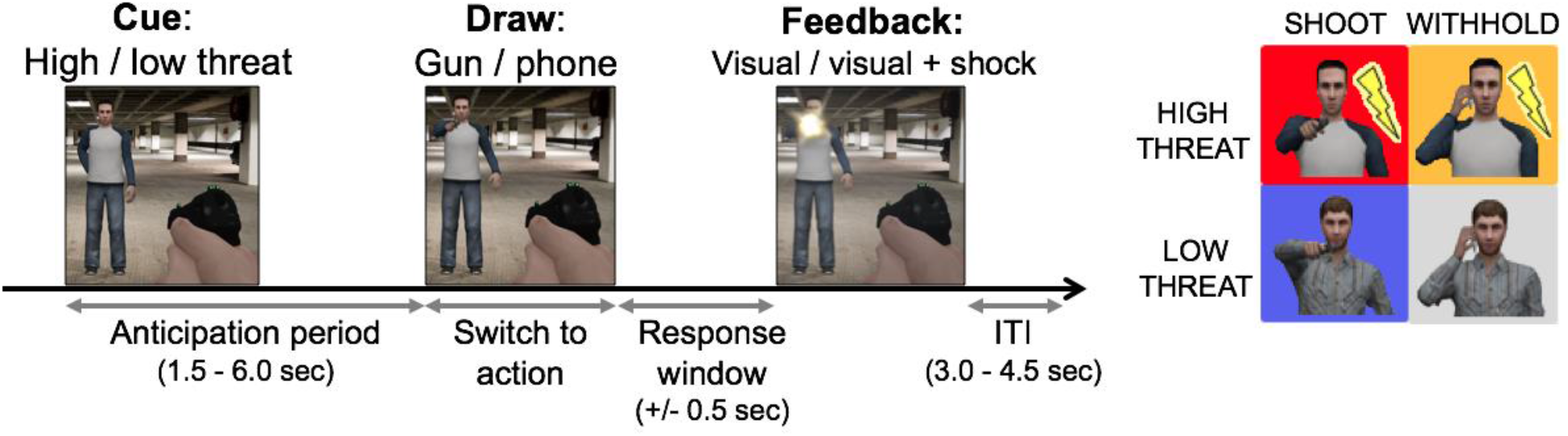
Schematic representation of the shooting task. One of two visually distinctive opponents is shown indexing either threat of shock (high threat cue) or shock safety (low threat cue) for a variable anticipation period of 1.5 to 6 seconds. After this, the opponent either draws a gun or a phone. A gun signals the participant to shoot (by button press) whereas a phone requires the participant to withhold responding. Incorrect shooting decisions (too slow, too fast, or false alarms) are indicated by visual feedback and on high threat trials (but not on low threat trials) followed by an electric shock.

Trials began with one of two visually distinctive opponents (the cue), standing in a parking garage, which signaled the preparation for potential action. One of the opponents was associated with threat of an aversive electrical shock (high threat cue) whereas the other signaled shock safety as it was never associated with an electrical shock (low threat cue). Which opponent was associated to the threat of shock was counterbalanced across participants and the order of high and low threat trials was randomized. After a short (500-1500ms), middle (1500-6000ms), or long (6000-6500ms) preparation period, the opponent either drew a gun (indexing a shooting response) or a mobile phone (indexing a withhold response). Armed opponents fired the gun after a brief delay forming a response window (approximately 500ms but individually titrated, see below) in which participants could shoot the opponent by a button press before being fired upon.

Participants firing too slowly were shot by the opponent, whereas participants firing too early (before the draw) or mistakenly (on withhold trials; false alarms) were shot by a policeman standing in the back of the garage. Only for the high threat opponent, these erroneous events (response that was too early or a false alarm) were followed by an aversive electric shock. To keep the frequency of aversive electric shocks consistent during the task and between subjects, the participant’s response window (time between the opponent drawing the gun and firing) was titrated in a way that participants would be shot on +/− 50% of the trials. This was accomplished by an algorithm that dynamically adjusted the duration of the response window (based on the participant’s median reaction time (RT) across low and high threat conditions) resulting in faster or slower firing actions of the opponent. Participants viewed their own “ in-task” hands, holding a gun during the entire trial and could fire at any time. Before the start of the task, electric shocks were set to an unpleasant but not painful level with a standardized work-up procedure which conisted of five electrical shocks of variable intensities that were rated on their unpleasentness (adapted from Klumpers et al., 2010).

Participants received explicit instructions on which opponent was associated with threat of shock and were verbally checked whether they understood the threat contingencies. To get acquainted with the task and threat contingencies, participants first underwent two blocks of 52 fast-paced training trials (80% short preparation intervals and shortened inter-trial intervals, ITIs). The final measurement phase consisted of three blocks of 28 trials including long anticipation intervals (80%) to allow for a sufficient number of trials for which the time course of body sway reductions and bradycardia could be analysed. Short and middle intervals were still presented for the rest of the trials to make the moment of attack unpredictable. ITIs varied between 3.0 to 4.5 seconds.

### Data collection and recording

#### Hair cortisol assessment

Hair strands with average lengths of 2.78 cm (range: 1 cm to 3 cm) were cut scalp-near from a posterior vertex position. These segments estimate the cortisol secretion over the last 3 months approximately given that the average hair growth rate is one cm per month. HCC was determined via a LC-MS/MS-based method (Gao et al., 2013) which is a selective and reliable procedure for the assessment of cortisol concentrations in hair samples. As hair sampling was restricted to participants with sufficient scalp hair length, HCC could be assessed for 343 participants. Six participants showed relatively high HCC values (above 3 standard deviations from the mean) and were therefore excluded from the analyses. We additionally used non-parametric tests to protect against the risk for false positives due to positive skewing observed in the remaining data (Rousselet & Pernet, 2012).

#### Trait anxiety

Before the shooting task, participants completed questionnaires including the Dutch version of the Spielberger Trait Anxiety Inventory (STAI), a widely used 20 item self-report instrument to assess general trait levels of anxiety (van der Ploeg, 1984). Internal consistency of trait anxiety scores (STAI-trait) was high as indicated by a Cronbach’s alpha of 0.89.

### Psychophysiological recording

#### Stabilometric platform

The task was performed on a custom-made stabilometric force platform (dimension 50 cm x 50 cm) that was located in front of a monitor displaying the task (see Figure 4a). The force plate consisted of four sensors that measured the displacement of the centre of pressure, or body sway, both in the anterior-posterior (AP) and medio-lateral (ML) direction. Participants were instructed to stand as still as possible and stood in a relatively stable position with feet approximately 30 cm apart.

#### Heart rate

Bipolar ECG was monitored using an ambulatory device (BrainAMP EXG MR 16 channel and EXG AUX apparatus) and three adhesive AG/AG-CL electrodes attached diagonal and above the heart. Raw ECG data was filtered with a Butterworth band-pass filter and was subsequently assessed via an in-house developed automatized R-peak detection algorithm. Peak detection was visually inspected trial-by-trial and corrected manually whenever required (e.g. when due to movement artefacts peaks could be not discerned). To assess the internal reliability of freezing measures including body sway and heart rate, we performed non-parametric Spearman’s rho correlations between odd and even trials. Analyses indicated a moderate to high reliability for body sway (low threat Rs = 0.62, p<0.001; high threat Rs = 0.78, p<0.001) as well as heart rate data (low threat Rs = 0.74, p<0.001; high threat Rs = 0.75, p<0.001).

### Data processing and statistical analyses

Raw electrocardiogram (ECG) and body sway data were recorded and downsampled to 125 Hz (with an initial sampling frequency of 2500Hz). The raw signal was filtered with a Butterworth band-pass filter (body sway: 0.01-10Hz (Demura & Kitabayashi, 2006), heart rate: 0.5-10 Hz (Laborde, Mosley, & Thayer, 2017; Butterworth, 1930).

### General task effects

#### Behavioural analysis

Processing and analyses of the behavioural data was carried out with Matlab 2015a and SPSS 23.0. RT analyses involved only correct responses within the response window of 200-500ms. A non-parametric Wilcoxon signed rank test was used to investigate differential effects of high and low threat trials on RT to minimize non-normality issues as well as outlier concerns. For our binary shooting accuracy measure (correct vs. incorrect) we performed Bayesian linear mixed models (Dixon, 2008; Jaeger, 2008; Quené & van den Bergh, 2008) in R using the brms package (Bürkner, 2016) which interfaces to Stan (Carpenter ea., 2016). The model included cue (high vs. low threat) and draw (shoot vs. withhold) as fixed factors and subject-specific effects as random-factors using a binomial distribution. Categorical factors were coded using sum-to-zero contrasts. The model included a random intercept per participant and random slopes for cue and draw and their interaction. To follow up the observed interaction effect between cue and draw, we modelled the cue effect (high vs. low threat) separately within the shooting as well as the withhold conditions. We fitted the models with 5 chains with 6000 iterations each (2000 warm-up). We used brms’ weakly informative default priors and a coefficient is reported as statistically significant when the associated 95% posterior credible interval is nonoverlapping with zero.

#### Psychophysiological analysis

Preprocessing was performed with Matlab2015a and statistical analyses were carried out in SPSS23. Analysis of body sway and heart rate included only trials with duration of minimally 6 seconds (Gladwin et al., 2016). As 80% of all trials had sufficiently long intervals to analyse body sway and heart rate, there were in total 28 trials per condition (high/low threat) for the analysis. The first trial from each block, and trials with insufficient signal-to-noise ratio, for example where R-peaks were non-detectable due to movement artefacts, were discarded from the analysis. A minimum of 50% remaining artefact-free trials (14 trials) for each condition were deemed necessary to include a participant’s dataset within the analysis (Hashemi et al., in press). For the heart rate analyses, six participants’ data were removed based on this criterion, and two additional participants were excluded because of incidental findings of arrhythmia. For the body sway analyses, two participants were removed due to hardware problems or discontinuation of the measurement due to dizziness.

In line with previous analyses of human freezing reaction, reductions in body sway in the AP direction were taken as an index of postural freezing (Gladwin et al., 2016; Roelofs et al., 2010). Due to the spaced feet position (30 cm apart) on the balance board, the AP direction has a larger movement range and therefore a greater sensitivity to affective modulations as well as individual differences in freezing compared to the ML direction (Hagenaars, Stins, & Roelofs, 2012; Niermann et al., 2017). Similarly, reductions in heart rate reflect bradycardia responses. During the anticipation interval, event-related changes were calculated between 2.5 to 6.0 seconds for body sway and heart rate (inter-beat interval, IBI), relative to a baseline period of 1 second before cue onset. The time window of the analysis was chosen to exclude non-specific orienting effects from threat-related prolonged bradycardia and body sway (Hagenaars et al., 2014; Hermans, Henckens, Roelofs, & Fernández, 2013). To test whether anticipation of threat was related to body sway reductions and enhanced bradycardia, we ran a non-parametric Wilcoxon signed-rank test comparing contrast values of high vs. low threat [high - low threat] during the anticipation window against the baseline period. A non-parametric test was chosen instead of ANOVA in order to minimize non-normality and outlier concerns.

### Freezing-reactions and their relation to subsequent actions

Previous studies pointed to a role of freezing in action preparation showing its adaptive value in active threat contexts (Gladwin et al., 2016; Hashemi et al., in press). To verify a similar relationship within the current study, we performed rank correlations (Spearman’s’ rho) between freezing measures (as assessed with body sway and bradycardia) and subsequent actions including RT and accuracy. To obtain a freezing index per individual, we averaged body sway as well as heart rate (IBIs) responses within the anticipation period (2.5s - 6.0s) for high and low threat trials separately, and computed the high-low threat contrast on these averages. However, as the previous studies showed similar associations between freezing and action preparation on high and low threat trials, we additionally tested specifically the high threat condition (rather than the difference scores) as action preparatory states are expected to be most pronounced here. To minimize chances of type-II errors due to the multiple testing (here six tests for each freezing-related measure), we corrected p-values using a false discovery rate (FDR) adjustment, which considers that the tested values might be interdependent, via the *p.adjust* function from the stats package (R Core Team, 2015).

### Individual difference analyses of freezing reactions

We performed non-parametric Spearman’s rho correlations (α < 0.05) to investigate our main aim of the study and test whether individual differences in threat-induced freezing (high - low threat contrast), as assessed with body sway and bradycardia, are associated with lower HCC and higher trait anxiety. P-values were again adjusted with FDR for two comparisons for each freezing measure. To control whether found associations are driven by the high threat condition as a consequence of fear of threat of shock we subsequently tested for the relationships within the high threat condition.

Given recent concerns about reproducibility of scientific findings (Ioannidis, 2005; Jasny, Chin, Chong, & Vignieri, 2011) and the small effect sizes that can be expected for individual differences analyses (Schönbrodt & Perugini, 2013), we additionally split all participants randomly into two independent groups matched on age, gender, and group (police recruits or controls) for split-sample validation (see Supporting Information). All results reported below remained significant in both sub-samples, except for one (see below) for which the effect size was smaller than expected (<0.2) and required the power of the full sample.

## Results

### General task effects

#### Threat of shock induced faster - but also more impulsive shooting

A significant cue [high vs. low threat] by draw [gun vs. phone] interaction was observed on accuracy (B= 0.67, 95% CI [0.42, 0.93]). High threat (compared to low threat or shock safety) was related to higher accuracy when required to shoot (B= −0.41, 95% CI [−0.7, −0.1]) but was also linked to more shooting errors when required to withhold responding (false alarms; (B= 0.34, 95% CI [0.25, 0.43]) see Figure 2a). Additionally, participants’ correct shooting actions were faster under high threat as compared to low threat (Wilcoxon p<0.0001, see Figure 2b). These results verify that our threat manipulation was successful as they are in line with previous findings showing that (high) threat is related to more impulsive shooting behaviour (Hashemi et al., in press; Nieuwenhuys et al., 2012).

**Figure 2.**
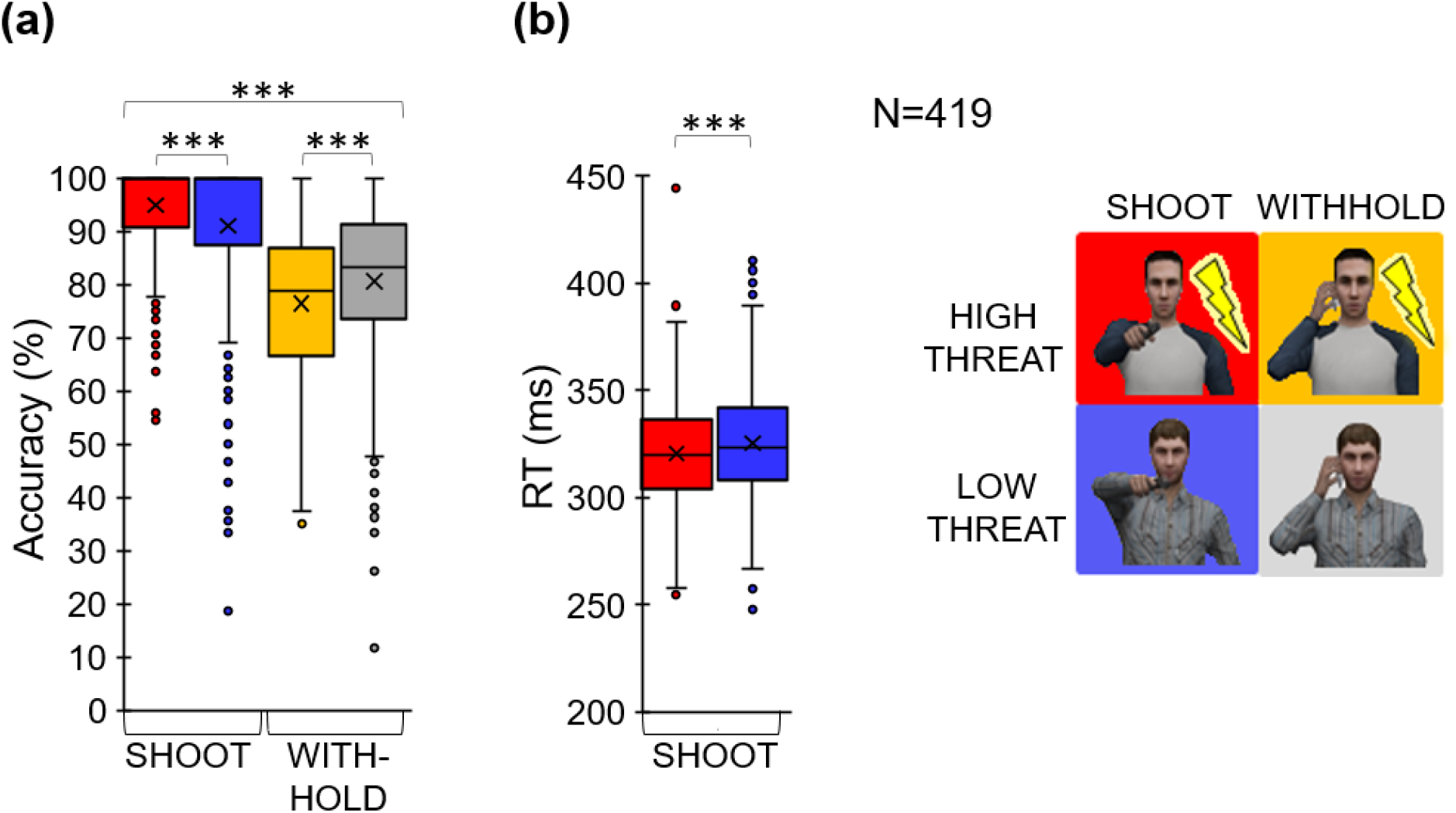
a) Across participants, anticipation of high threat was followed by higher accuracy when required to shoot, but also more errors when required to withhold shooting. b) Correct shooting responses were on average faster for high compared to low threat trials across participants. Boxplots illustrate the data distribution, with the middle line indicating the median and the edges of the box indicating 1^st^ and 3^rd^ quartiles. The mean is indicated by a cross. *** = p<0.001.

#### Anticipation of threat of shock increased body sway and heart rate reductions

Body sway and heart rate were generally decreased after cue onset (Wilcoxon: baseline vs. anticipation: body sway: p<0.001, 95% CI of the difference [4.37, 5.01]; heart rate: p<0.001, 95% CI of the difference [0.69, 0.81]), which is in line with previous studies that show these patterns during action preparation also in the relative absence of threat (e.g. Gladwin et al., 2016; Jennings & Van Der Molen, 2005). Replicating previous work (Hashemi et al., in press), reductions in body sway after cue onset were however stronger for high threat (threat of shock) compared to low threat (shock safety) (Wilcoxon [high – low threat]: baseline vs. anticipation] p<0.001, r= 0.1, 95% CI of the difference [0.04, 0.12], see Figure 3). Similarly, anticipatory heart rate decelerations were increased under high compared to low threat (Wilcoxon [high – low threat]: baseline vs. anticipation] p<0.001, r= 0.13, 95% CI of the difference [0.47, 0.85], see Figure 3). Consistent with previous findings (Azevedo et al., 2005; Niermann, Figner, Tyborowska, Cillessen, & Roelofs, 2018; Roelofs et al., 2010), body sway reductions and bradycardia were positively correlated (high threat: Rs=0.26 p<0.001, low threat: Rs=0.24 p<0.001 see Supporting Figure 3). Thus, body sway reductions as well as bradycardia during anticipation were magnified by the level of threat and the strength of these reactions was positively associated, indicative of human freezing reactions.

**Figure 3.**
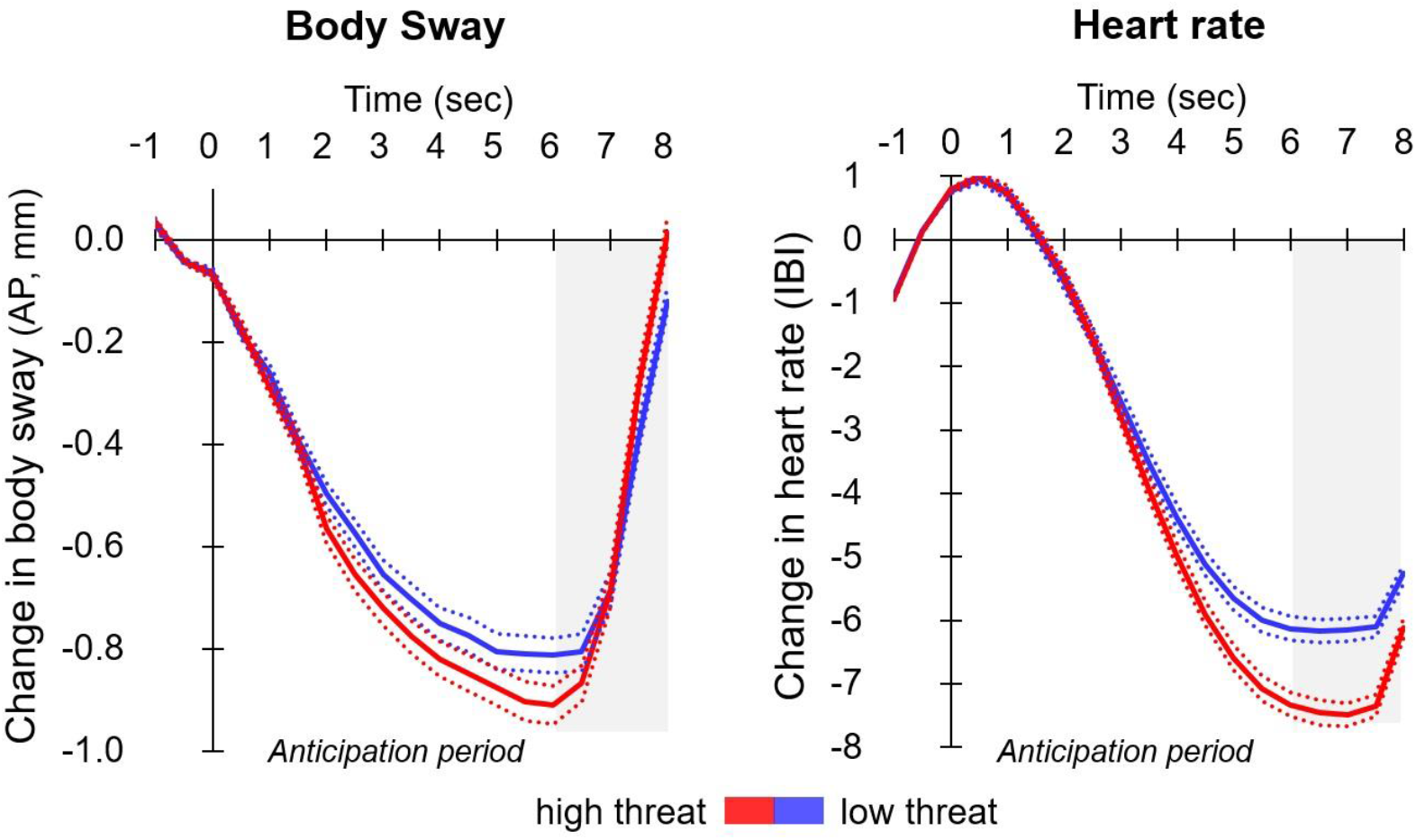
Anticipation of high threat (threat of shock) was associated with reduced body sway as well as bradycardia (compared to low threat trials) indicative of human freezing reactions. This is shown in stronger mean body sway reductions (left) during high threat, assessed in anterior-posterior (AP) deviations of centre of pressure, as well as enhanced mean heart rate decelerations (right), reflected in changes in inter-beat intervals (IBI). Standard errors of the mean are illustrated in dotted lines.

#### Anticipatory body sway and bradycardia are linked to the speed and accuracy of subsequent defensive action

Stronger reductions in body sway as well as heart rate were associated with faster subsequent shooting under high threat (body sway [high threat]: Rs = 0.17 p<0.001, [high-low threat]: Rs=0.07, p=0.18; heart rate [high threat]: Rs= 0.21 p<0.001, [high-low threat]: Rs= 0.14, p=0.015, all FDR-corrected, see Figure 4). Reductions in body sway, but not in heart rate, under threat of shock were also associated with more false alarms -erroneous shooting-when required to withhold responding (body sway: [high threat]: Rs= 0.16 p<0.003, [high-low threat]: Rs= −0.09 p=0.1; heart rate: ([high threat]: Rs= 0.03, p=0.571, [high-low threat]: Rs= 0.04 p=0.48, all FDR-corrected, see Figure 4). A post-hoc comparison of the correlation coefficients for body sway and heart rate confirmed a significantly larger association to false alarms for body sway (Fischer’s z = 1.88 p=0.03). These results indicate a robust link (see Supporting Information) between freezing-related bodily responses and subsequent actions (Gladwin et al., 2016; Hashemi et al., in press) but also suggest distinct predictive values for heart rate and body sway measures.

**Figure 4.**
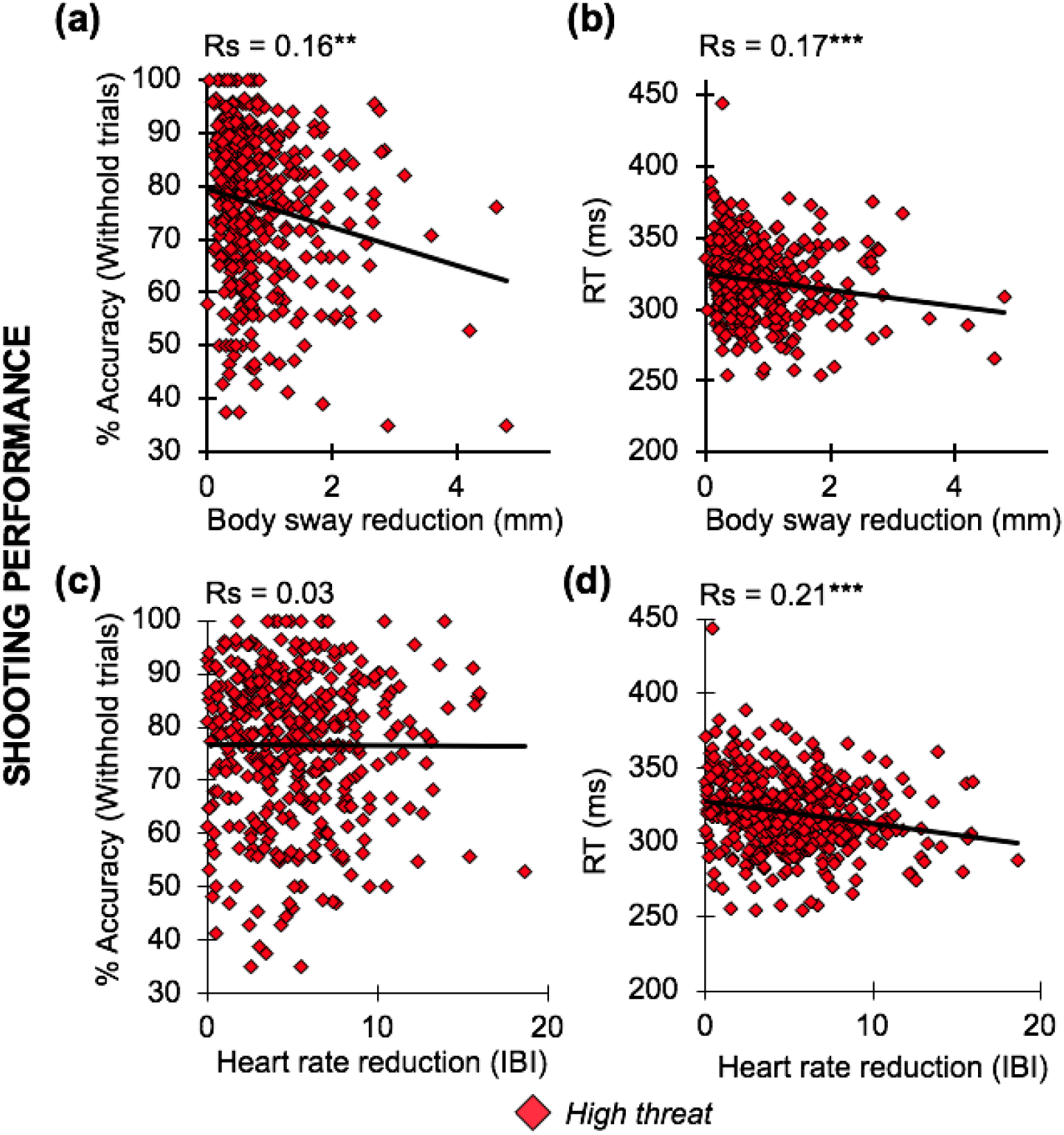
During the anticipation of high threat (threat of shock), mean body sway reductions were associated with more subsequent false alarms when required to withhold (a) as well as faster correct mean reaction times when shooting was required (b). Anticipatory heart rate decelerations under high threat were associated with faster correct mean reaction times (d) but not with accuracy on withhold trials (c). For illustrative purpose, absolute values of body sway and heart rate reductions are plotted (each point represents a participant). **= p<0.01, ***= p<0.001 FDR-corrected p-value for multiple comparisons.

### Individual difference analyses of freezing reactions

#### Freezing-related body sway reduction is linked to low HCC and high trait anxiety

##### HCC

On average, HCCs were 9.74 pg/mg (SD=16.44) which is in line with previously reported healthy populations (Stalder & Kirschbaum, 2012). As predicted, stronger threat-enhanced body sway reductions (high – low threat) was associated with lower HCC (Rs= 0.13, p=0.028 N=341, FDR-corrected, see Figure 5). A post-hoc analysis indicated that this association was driven by the expected association under high threat (R=0.11, p=0.052 FDR-corrected). As recent reviews on hair cortisol assessments suggest that HCC covaries with age, gender, hair washing frequency as well as hair treatment (Greff et al., 2018; Stalder et al., 2017), we ran additional partial correlation analyses to control for all these confounding variables. The association between HCC and body sway reductions became slightly more robust in this control analysis (R=0.15; p=0.009). Note that the effect sizes are small according to conventional criteria but effect sizes commonly decrease with increasing sample sizes (Schönbrodt & Perugini, 2013). HCC did not correlate with threat-enhanced changes in bradycardia (high – low threat) (Rs= −0.07, p=0.21, N=337, FDR-corrected) or shooting performance (RT and accuracy, lowest p-value of all: p=0.2 FDR-corrected) and also not after controlling for possible confounds.

**Figure 5.**
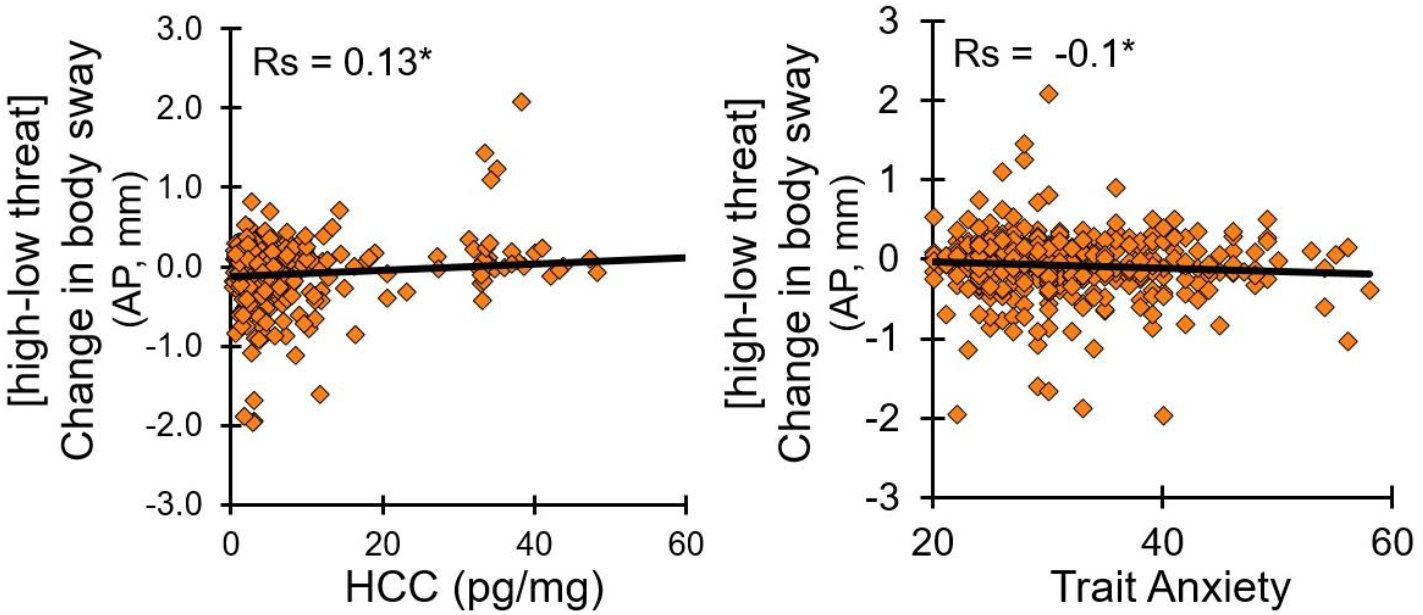
Stronger threat-related reduction in mean body sway (high-low threat) were linked to (left) lower HCC and (right) higher trait anxiety across participants. Spearman’s’ rank (Rs) correlations were performed to minimize non-normality and outlier concerns. * = p<0.05 FDR-corrected for multiple comparisons.

##### Trait anxiety

Stronger threat-enhanced body sway reductions (high – low threat) show also a small negative association with trait anxiety (Rs= −0.10, p=0.04 N=417, FDR-corrected, see Figure 5). Again, control analysis indicated that this effect was driven by high threat (Rs= −0.10, p=0.052 FDR-corrected). There was no such association with trait anxiety for threat-enhanced changes in bradycardia (high – low threat) (Rs= 0.01, p=0.852, N= 411, FDR-corrected). Post-hoc analyses did not show an association of HCC and trait anxiety within our sample (Rs= −0.06 p=0.26 N=343), also not when controlling for age, gender, hair washing frequency, and hair treatment (R=0.04, p=0.48). No other factors, including shooting performance (RT and accuracy, lowest p-value of all: p=0.43 FDR-corrected) were related to trait anxiety.

All results reported above are cross-validated in our split sample analysis (that is they remain significant in each of our randomly generated splits of the dataset, see Supporting Information for details). Only the link between trait anxiety and threat-related body sway reductions (high – low threat), showed a similar direction in both samples but reached significance only in one sub-sample (Sample 1 Rs= −0.08, p=0.23 (N=208); Sample 2 Rs= −0.14, p=0.05 (N=209)).

## Discussion

The aim of this study was to reveal whether individual differences in human freezing reactions relate to subsequent defensive reactions and two key predictors of stress-related psychopathology: HCC and trait anxiety. After verifying that our threat manipulation produced the expected freezing effects, we showed that body sway - as well as heart rate reductions - were robustly associated with faster accurate shooting which supports the view of the adaptive nature of freezing-related reactions within an active task context. Critically, our findings however also demonstrate that individual differences in threat-induced body sway reductions, but not bradycardia, were robustly related to 1) lower accuracy when required to withhold shooting, 2) lower accumulated HCC, and 3) higher trait anxiety. Together, these results demonstrate for the first time in a well-powered human sample that postural freezing in humans may relate to stress-relevant features of coping and threat processing.

Previous studies on the relation between freezing and acutely circulating basal cortisol levels predominantly indicated a positive association (Kalin et al., 1998; Niermann et al., 2017). Here we focussed on long term accumulating cortisol levels and, in line with work on hair cortisol by Hamel et al. (2017) in primates, we find that lower HCC is associated with more postural freezing. These contrasting findings are likely due to the different time scales of these measures, with acutely circulating cortisol levels reflected in saliva, plasma, or urine versus accumulated cortisol concentrations of several months in hair. As such, HCC can be viewed as a long-term, more state-independent marker of HPS-axis functioning. Similar opposing results were reported for anxiety as well as PTSD where acute cortisol assessments from saliva were positively related to anxiety and PTSD (Mantella et al., 2008; Vreeburg, Zitman, Van Perl, & Al., 2010), whereas chronic cortisol levels assessed from hair were negatively related to anxiety and PTSD (Steudte-Schmiedgen et al., 2015; Steudte et al., 2011). This contrast highlights the added value of investigating the link between freezing-related reactions and cortisol signalling with long-term, accumulated measures as in hair.

In addition to the association to HCC, threat-related postural freezing was also linked to higher trait anxiety. This result is in line with previous work indicating prolonged freezing reactions during anxiety (Frank et al., 2006; Lopes et al., 2009; Niermann et al., 2017; Roelofs et al., 2010). For example, during a passive viewing task, Roelofs et al. (2010) showed that enhanced freezing in response to threatening pictures was related to state anxiety in healthy participants. Previous associations were based on relatively small samples and passive tasks where active threat coping was not an option. We therefore present the first evidence that individual differences of anticipatory freezing reactions are linked to trait anxiety, also in conditions where timely and accurate responding could have prevented threat of shock.

The link between stronger freezing with subsequent threat coping, lower hair cortisol as well as higher trait anxiety raises the question whether threat-potentiated freezing reactions reflect an adaptive or maladaptive reaction to threat. In line with previous research (Hashemi et al., in press), we observed that both (heart rate and postural) measures of freezing were associated with faster action decision-making. However, given that reduced body sway was uniquely associated with both predictors of stress vulnerability and increased error rates, it may well be that excessive (here postural) freezing also constitutes a somatic marker of stress vulnerability. To address this question, it is helpful to consider patterns of freezing in vulnerable individuals with stress-related symptoms. In non-clinical healthy samples, experiencing aversive life-events as well as anxiety were associated with *increased* freezing reactions (including both reductions in body sway and heart rate, Hagenaars et al., 2012; Roelofs et al., 2010). In contrast, clinical samples of PTSD patients were shown to express *reduced* freezing reactions (Adenauer, Catani, Keil, Aichinger, & Neuner, 2010; Fragkaki, Roelofs, Stins, Jongedijk, & Hagenaars, 2017; Orr & Roth, 2000). This is thought to result from increased hyperarousal and excessive sympathetic reactivity that supresses the adaptive, preparatory parasympathetically-dominated freezing reaction. By this, the hierarchically-organized defence cascade that normally shifts from initial preparatory parasympathetically-dominated freezing to immediate sympathetically-dominated fight-or-flight actions under increasing and upcoming threat, seems to be disrupted (Bradley, Codispoti, Cuthbert, & Lang, 2001; Lang, Davis, & Öhman, 2000; Mobbs, Hagan, Dalgleish, Silston, & Prévost, 2015). Accordingly, one could argue that in our healthy sample, increased freezing reactions may signal heightened responsivity to threat that may be still adaptive. However, due to the relation between postural freezing and predictors of stress vulnerability (low HPA-axis activity and anxiety), increased postural freezing may reflect a heightened vulnerability to future challenging or traumatic experiences. As such, we speculate that traumatic life-experiences would turn this heightened threat responsivity into immediate sympathetic activations, which may lead to reduced freezing-related reactions as described previously in PTSD (Adenauer et al., 2010). Future longitudinal investigations are needed to address this issue within a prospective framework as well as in clinical samples to explore whether threat-enhanced freezing and low HPA-axis activity is linked to resilient or maladaptive stress responding.

We note some limitations and interpretational issues. First, it is a strength that the association between body sway and HCC was replicated in our cross-validation approach. The relationship between body sway reductions and trait anxiety was however not replicated in one of our cross-validation samples which indicated a potentially similar effect but insufficient power. A related point is that the effect size of the observed correlations is small by traditional standards (i.e., coefficient between 0.1-0.15). Recent meta-analyses however show that traditional guidelines for interpreting correlation coefficients may have been too stringent (Gignac & Szodorai, 2016; Hemphill, 2003), potentially because observed correlations between experimental measures are always dampened by the imperfect measurement reliability of the variables (Hedge, Powell, & Sumner, 2017a; Vul, Harris, Winkielman, & Pashler, 2009). Furthermore, evidence from simulations show that increasing sample size is generally associated with decreasing correlation coefficients and that a large sample size (e.g. N>250) more accurately captures the true effect size (Schönbrodt & Perugini, 2013). The small effect size of the correlations found within our large sample therefore suggests we were able to capture a robust underlying association that is still of theoretical significance. Finally, all associations found in relation to the HPA-axis activity and anxiety were specific to body sway reductions, as opposed to concomitant bradycardia responses, which are in some studies taken as proxy for freezing when measurements of body sway are not possible (e.g. neuroimaging environments Hashemi et al., in press; Wendt, Löw, Weymar, Lotze, & Hamm, 2017). Our results may point to the specificity as well as sensitivity of body sway reductions (within an active task context) to factors that are associated with stress vulnerability. Previous associations found between individual differences in stress vulnerability and freezing reactions were not consistently associated with measures of only body sway or heart rate (Hagenaars et al., 2012; Niermann et al., 2017). This variability could be due to task demands (active vs. passive tasks) or subject-specific factors. Bradycardia reactions have been often studied in the context of (threat) anticipation processes that are associated with optimized perceptual and motor processes for subsequent actions (Lojowska et al., 2015; Löw, Lang, Smith, & Bradley, 2008; Löw et al., 2015; Porges, 2003). Our result of an association between bradycardia and reaction times is therefore in line with this literature. Note that this is distinct from studies investigating tonic (e.g. not stimulus-locked) changes in heart rate variability (HRV), which generally show relations with maladaptive responding and increased stress vulnerability (Beauchaine & Thayer, 2015; Lyonfields, Borkovec, & Thayer, 1995). Given the consistent relationship found between heart rate and body sway reductions during threat anticipatory freezing reactions (Niermann et al., 2017; Roelofs et al., 2010) and the fact that both measures were found to have a comparable measurement reliability in our study, it would be interesting for future studies to reveal in how far differential effects between body sway and bradycardia reactions can be explained.

Concluding, individual differences in postural freezing as assessed with body sway reductions were related to subsequent defensive reactions as well as stress vulnerability indexed by lower HCC and higher trait anxiety. This implies that somatic reactions such as postural freezing may help prime defensive actions and are potentially mechanistically related to the HPA-axis changes and maladaptive threat processing that have been implicated in anxiety disorders.

## Author information

### Contributions

This work was supported by a VICI grant (#453-12-001) from the Netherlands Organization for Scientific Research (NWO) and a starting grant from the European Research Council (ERC_StG2012_313749) awarded to KR, also supporting MH, WZ, RK, FK and SBJK. Study concept and experimental design was done by MMH, FK, KR. MMH, R.J., WZ, RK and SBJK contributed to data acquisition. Analysis was performed by MMH, supported by FK, and BF helped with analysis for the Bayesian models. The manuscript was drafted by MMH, KR and FK and all co-authors reviewed and gave feedback on the manuscript. The authors gratefully acknowledge Annika Smit and Vanessa van Ast for their help with initiating the study and recruitment as well as all research assistants and master students for their help with data acquisition.

### Data availability

The datasets analysed during the current study are available from the corresponding author upon reasonable request.

### Code availability

The code that has resulted in the reported findings is available from the corresponding author upon reasonable request.

### Corresponding author

Correspondence to M. M. Hashemi (, +31639126363) and K. Roelofs

