## Supporting Information for "Human defensive freezing is associated with acute threat coping, long term hair cortisol levels and trait anxiety"

### Methods

To confirm the reliability of our results given the reproducibility crisis in science (Ioannidis, 2005; Munafò et al., 2017), we randomly split the sample into two groups matched on age, gender, and group (police recruits or healthy civilians) for cross-validation. Previous work on human freezing reactions showed a link between freezing-related periaqueductal brain activation and action preparation with medium effect size (R=0.37, N=54, Hashemi ea. in prep). As the sample size of this former study was relatively small for individual differences analysis, the effect size may have been inflated (Hedge, Powell, & Sumner, 2017). Given a robust but imperfect expected test-retest reliability of our measures (Niermann, Figner, Tyborowska, Cillessen, & Roelofs, 2018), we expected effects of interest to be rather small (H1: r= 0.2). For such a small effect size, power calculations estimated a minimum sample size of 207 participants including the following parameters $\alpha$ <0.1, 1-β= 0.95, H0: r=0 (Faul, Erdfelder, Lang, & Buchner, 2007), which could be approximated with our two samples on most analyses (see descriptive details in Table 1).

| | Full Sample | Sample 1 | Sample 2 | Group difference (p-value) |
|---|---|---|---|---|
| *Demographics* | | | | |
| Age (M, SD) | 24.17 (5.1) | 24.09 (5.18) | 24.24 (4.99) | p=0.48 |
| Male (N) | 313 | 157 | 156 | |
| Police (N) | 337 | 169 | 168 | |
| *Psychophysiological measures* | | | | |
| Body sway (mm) [threat-safe] | -0.08 (0.39) | -0.11 (0.38) | -0.05 (0.39) | p<0.05* |
| Heart rate (bpm) [threat-safe] | -0.75 (2.1) | -0.55 (2.13) | -0.95 (2.05) | p=0.09 |
| *Hair-related variables* | | | | |
| HCC (pg / mg) | 9.74 (16.44) | 10.1 (15.81) | 9.4 (17.07) | p=0.16 |
| Washes per week (M, SD) | | 3.53 (1.22) | 3.65 (1.11) | |
| Hair color (%) | | | | |
| ▪ blond | | 46.43 | 43.43 | |
| ▪ brown | | 41.67 | 45.71 | |
| ▪ red | | 4.17 | 3.43 | |
| ▪ black | | 7.74 | 6.86 | |
| ▪ grey | | 0 | 0.57 | |
| *Psychological traits* | | | | |
| Trait Anxiety (STAI) | 31.31 (7.4) | 30.89 (6.85) | 31.73 (7.91) | p=0.43 |

**Table 1.** Demographics, psychophysiological measures, hair-related variables, and psychological traits are summarized for the full sample, Sample 1 and Sample 2. A non-parametric Mann-Whitney U test was performed to test for differences between Sample 1 and Sample 2. Sample 1 showed stronger mean threat-related body sway reductions compared to Sample 2. No other sample-related differences were observed. HCC= hair cortisol concentrations.

### Results

#### Defensive freezing-related reactions link to subsequent actions

Stronger body sway reductions were associated with faster correct shooting ([high threat]: Sample 1 Rs = 0.2 p<0.01 (N=208); Sample 2 Rs = 0.14 p<0.05 (N=209); see SI Figure 1b), but also more erroneous shooting when required to withhold responding under high threat ([high threat]: Sample 1: Rs= 0.18 p<0.05; Sample 2: Rs=0.15 p<0.05, see SI Figure 1a). Heart rate responses were similarly associated with faster correct reaction times under high threat (Sample 1: Rs= 0.16 p<0.05 (N=202); Sample 2: Rs=0.25 p<0.001 (N=209) see SI Figure 1d). In contrast to body sway reductions, heart rate decelerations showed no relationship to shooting errors when required to withhold responding under high threat (false alarms, Sample 1: Rs= -0.01, p=0.84; Sample 2: Rs=0.07 p=0.34 see SI Figure 1c). Together these results indicate that anticipatory freezing-related reactions link to subsequent shooting actions.

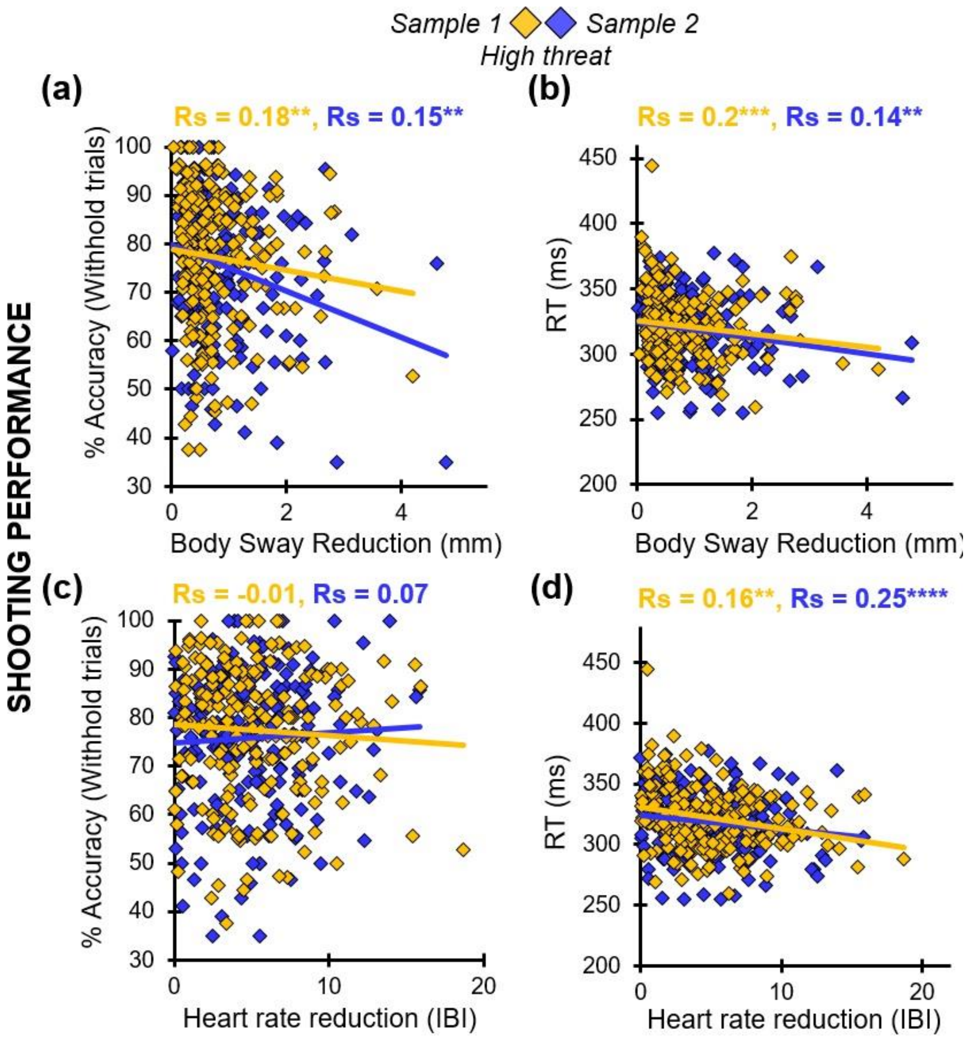


**Supplementary figure 1.** Stronger body sway reductions during anticipation of high threat (threat of shock) were robustly associated with (a) more erroneous shooting decisions when required to withhold as well as (false alarms) (b) faster reaction times when a response was required. Heart rate reductions under anticipation of high threat were (c) not related to false alarms (d) but were associated with faster reaction times. Rs = Spearmans Rho ** = p<0.05, *** = p<0.01, **** = p<0.001

**Individual difference analyses of freezing reactions**

**Body sway reductions relate positively to HCC and negatively to trait anxiety.**

**HCC.** Our cross-validation approach was not ideal as hair sampling was restricted to 343 participants resulting in two samples that were under the estimated sample size that are required to find small effects (N=207). Nevertheless, stronger threat-related body sway reductions during anticipation were associated with lower HCC ([high - low threat]: Sample 1 $Rs=0.15$, $p<0.01$ (N=167); Sample 2 $Rs=0.14$, $p<0.01$ (N=174), see SI Figure 2a). HCC was not correlated with threat-related bradycardia responses ([high - low threat]: Sample 1 $Rs= -0.02$, $p=0.8$ (N=163); Sample 2 $Rs= -0.12$, $p=0.11$ (N=174) see SI Figure 2c).

**Trait anxiety (STAI).** Threat-related, anticipatory body sway reductions showed a significant link to trait anxiety only in one sample ([high – low threat]: Sample 1 $Rs= -0.08$, $p=0.23$ (N=208); Sample 2 $Rs= -0.14$, $p<0.05$ (N=209) see SI Figure 2b). There were no other relationships observed, including bradycardia responses ([high - low threat]: Sample 1 $Rs= 0.05$, $p=0.46$ (N=202); Sample 2 $Rs= -0.03$, $p=0.7$ (N=209) see SI Figure 2d). Notice that the actual effect sizes are smaller than expected ($r=0.2$), and being able to detect such small effect sizes requires the power of the full sample. Together, these results indicate that threat-related postural freezing robustly predicts lower HCC, as well as higher trait anxiety although the latter association required the power of the full sample. This again highlights the relation of postural freezing with predictors of stress-related psychology.

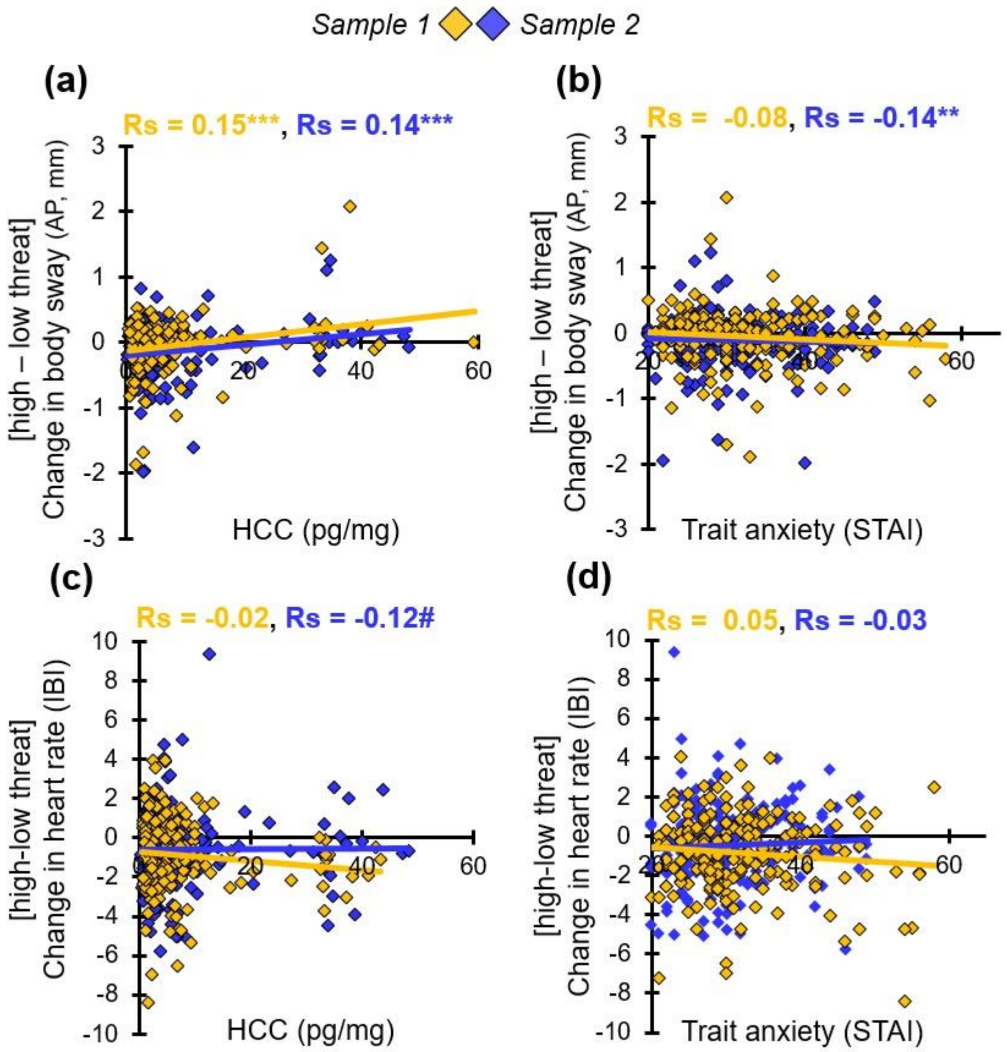


**Supplementary figure 2.** Individual differences in changes of threat-related body sway reductions (high-low threat) from baseline during anticipation robustly predicted (a) lower HCC. It also showed an association with (b) higher trait anxiety in one sample. Similar associations were not found in changes in heart rate (IBI) during anticipation, not for (c) HCC, nor for (d) trait anxiety. Spearmans' rank (Rs) correlations were performed to minimize non-normality and outlier concerns. ** = p<0.05, *** = p<0.01, # = p<0.11

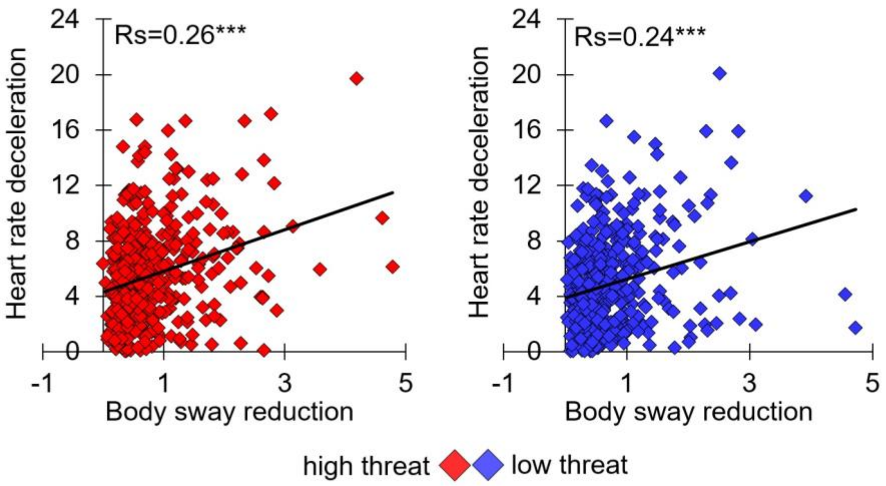


**Supplementary figure 3.** During anticipation of high and low threat the magnitude of body sway as well as heart rate reductions were correlated. ***p<0.001.
